# Fine needle aspiration biopsy of breast specimens effectively harvests cells for patient-derived organoids modeling breast ductal carcinoma in situ

**DOI:** 10.1101/2025.08.12.669895

**Authors:** Julia Ye, Nadine Goldhammer, Poonam Vohra, Shruti Warhadpande, George C. De Castro, Cristian Maldonado Rodas, Mark M. Moasser, Kirithiga Ramalingam, Shoko Abe, Michael Alvarado, Cheryl Ewing, Karen Goodwin, Rita Mukhtar, Jasmine Wong, Laura Esserman, Ronald Balassanian, Jennifer M. Rosenbluth

## Abstract

**Background:** Patient-derived organoids (PDOs) generated from benign breast tissue and breast carcinomas have successfully recapitulated their respective in vivo counterparts. PDOs model tumorigenesis and allow for screening of novel therapeutics personalized to individual patients. However, acquiring cells to generate PDOs is cumbersome. We demonstrate the feasibility of fine needle aspiration biopsy (FNAB) for harvesting cells for PDOs modeling ductal carcinoma in situ (DCIS).

**Methods:** Surgical specimens from patients with biopsy-proven DCIS were used for this study. Core needle biopsy (CNB) was performed on fresh specimens in the operating room, and tissue was mechanically dissociated before culture in basement membrane extract (BME) and organoid medium to generate PDOs. FNAB was performed in the gross room on fresh specimens, and the remaining aspirate was similarly submitted for PDO culture.

**Results:** PDOs were successfully generated in 15/18 specimens obtained by CNB and 7/11 specimens obtained by FNAB. The average time to initial organoid growth was 4 days for FNAB specimens compared to 19.3 days for CNB specimens. Tumor cells were seen on 7/11 FNAB smears and 16/18 CNB touch preps. Immunofluorescence staining confirmed the presence of both luminal and myoepithelial cells in derived PDOs.

**Conclusions:** FNAB effectively obtains cells for PDOs modeling DCIS. CNB after mincing yielded PDOs with a high success rate, but they were slow to establish. Notably, the time to organoid growth was significantly shorter for FNAB specimens. Thus, FNAB offers an efficient alternative for breast PDO culture and can reduce the time and resources spent on generating PDO cultures.

## Introduction

Organoids are in vitro cultures of various cell types within a 3-dimensional extracellular matrix and can be generated from primary explants of tissue fragments, clonal derivatives of primary epithelial stem cells, and epithelial-mesenchymal co-cultures from embryonic stem cells or induced pluripotent stem cells. [1,2] In the realm of cancer biology, patient-derived organoids (PDOs) derived from normal or diseased patient tissue are quickly becoming the new standard for basic and translational research, because they are able to capture multiple cell lineages and because the 3-D platform allows for cell-cell and cell-stromal interactions that better mimic the native tissue microenvironment. [3] Moreover, the ability of PDOs to capture the diversity of individual patient tissues allows for the improved study of cancer heterogeneity, as well as the screening for effective novel therapeutics and the investigation of predictors of therapeutic response and resistance in a patient-centered manner.

Breast cancer remains the most common cancer diagnosed in women in the United States and the second most common cause of cancer-related death in women in the United States, [4] despite rigorous screening guidelines and a multitude of available therapeutics. These statistics highlight the underlying disease heterogeneity of breast cancer and the need for improved models for breast cancer tumorigenesis. PDOs have been successfully produced from invasive breast carcinomas and ductal carcinoma in situ (DCIS), [5] as well as from benign breast tissue. [6] Importantly, these breast PDOs are able to be propagated long-term and have been shown to successfully recapitulate their respective in vivo counterparts in terms of histomorphology, cellular composition, and biomarker status.

However, specimen acquisition and initial PDO establishment can be labor-intensive processes that require significant resources, due to slow tissue growth and other factors that delay verification and validation procedures. In the context of breast PDOs, these issues can be exacerbated for in vitro models of precursor or high-risk lesions such as DCIS, where the need for detailed clinical evaluation of the target lesion often precludes anything other than minimal sampling of large lesions. At the same time, improved models of DCIS are needed to test cancer prevention or interception strategies that could spare patients morbidity and mortality from the development of invasive disease, given the heterogeneity of DCIS and that only 20-40% of DCIS cases progressive to invasive carcinoma. [7,8]

Standard protocols for breast PDO generation rely on microdissection or core needle biopsy (CNB) of excision specimens, followed by manual tissue mincing and/or enzymatic digestion. [5,9] Fine needle aspiration biopsy (FNAB) is an economical and minimally invasive biopsy technique that uses a thin beveled needle (typically 22-25 gauge) to sample both palpable and deep-seated lesions in patients. FNAB can also be performed on fresh surgical specimens at the bench. [10] Recent work has demonstrated that FNAB can successfully harvest cells to produce PDOs modeling melanoma, gastrointestinal carcinomas, thyroid carcinomas, and renal cell carcinomas. [11,12] Here, we examine the feasibility of using FNAB to harvest cells for PDOs modeling breast DCIS. In addition, we compare the efficacy of PDO generation from tissues sampled by FNAB compared to CNB.

## Methods

### Cell and tissue isolation from breast samples by CNB and FNAB for organoid generation

Specimens were collected from patients undergoing breast surgery who consented to donate tissue for research at the University of California, San Francisco (UCSF), under a protocol reviewed and approved by the Institutional Review Board. Total and partial mastectomy specimens from patients with biopsy-proven DCIS were sampled fresh by either CNB or FNAB, as shown in Figures 1–2.

**Figure 1.**
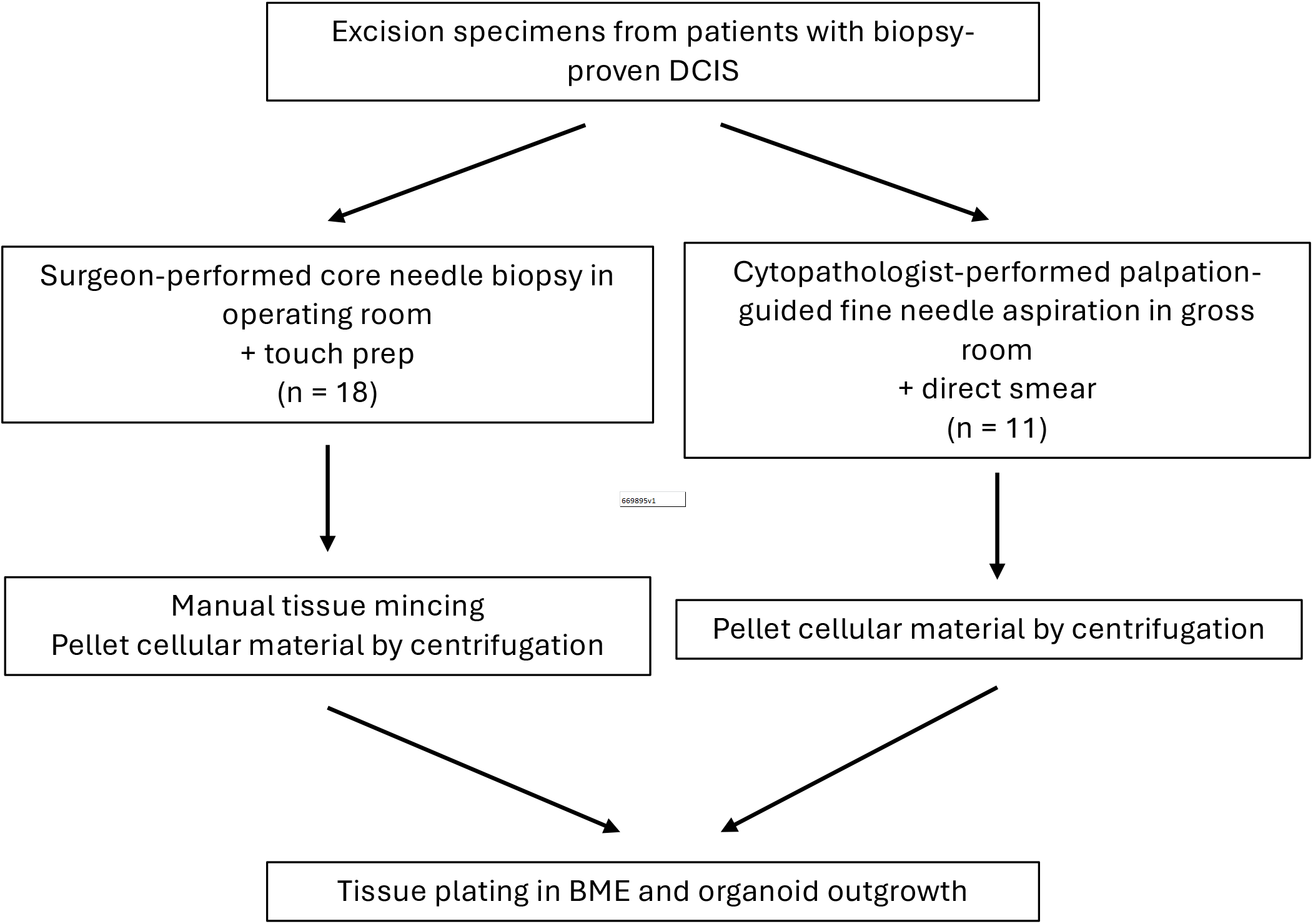
Study design.

**Figure 2.**
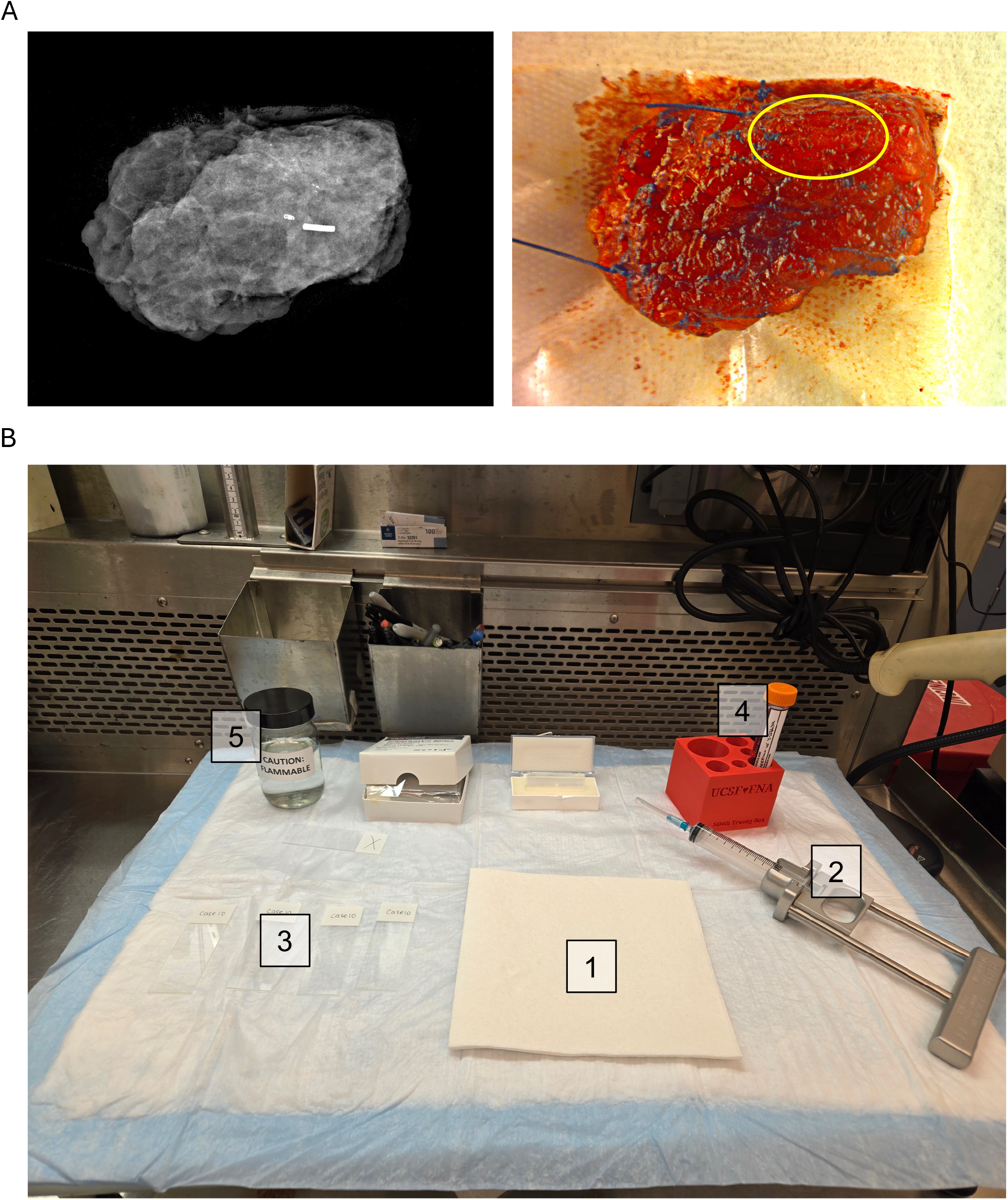
Palpation-guided FNAB of breast excision specimens. (A) Representative image of x-ray and gross photograph of a partial mastectomy specimen. Yellow circle corresponds to the area of calcifications as identified on the x-ray and indicates the area to be targeted by palpation-guided FNAB. (B) Bench set-up for palpation-guided FNAB with (1) clean paper towel and absorbency pad, (2) 10 ml syringe with 1 inch 23-gauge needle in an FNA syringe holder, (3) clean labeled slides for direct smears, (4) 15 ml tube of base medium with ROCK inhibitor, and (5) glass jar of 95% ethanol for direct smears.

CNB was performed by surgeons in the operating room in 3-4 passes with an 18-gauge spring loaded biopsy tool (BARD) after specimen x-ray to guide the site for the biopsy, and tissue was submitted in 0.9% normal saline or viably frozen. Touch preps were performed immediately prior to organoid culture for assessment by a breast pathologist and cytopathologist (see below under “Organoid culture”).

FNAB was performed by cytopathologists in the gross room after performing a specimen x-ray to identify the presence and location of calcifications, biopsy clips, and localization seeds. Palpation-guided FNAB was performed in 1-3 passes with 23-gauge beveled needles targeting the identified areas of interest. Direct smears were prepared from each pass to assess cellularity and adequate sampling of the targeted lesion. The smears were fixed in 95% ethanol and initially stained with Toluidine blue with a temporary coverslip. After microscopic review, the slides were returned to 95% ethanol, which removed the coverslip and allowed for the Toluidine blue stain to wash off. The smears were then permanently stained with Papanicolaou stain according to standard protocols. The remaining aspirate was submitted in base medium with ROCK inhibitor [Advanced DMEM/F12 (Fisher Scientific, BW12-719F) with 50 μg/ml Primocin (InvivoGen, ant-pm-05), 1x GlutaMax (Thermo Fisher, 35-050-061), 100 units/ml Penicillin/Stroptomycin (Thermo Fisher, 15140122), 10 mM HEPES (Thermo Fisher, 15-630-080), and 5 μM ROCK inhibitor Y-27632 (abcam, ab143784)].

### Organoid culture

Organoids were established and cultured as previously described. [5,6,9] A 2-3 mm^3^ piece of tissue was cut from each tip of CNB for formalin-fixed paraffin embedding (FFPE), and touch preps were performed with the remaining tissue. Touch preps were stained with Giemsa stain according to standard protocols. CNBs were then minced into 1 mm^3^ pieces and embedded in growth factor-reduced Cultrex Basement Membrane Extract (BME) type 2 (BioTechne, 3532-010-02). FNAB specimens were centrifuged at 300 x g for 3 min and the pellet was embedded in BME. Both CNB fragments and FNAB specimens were plated in 24-well plates and incubated at 37 °C until solidification of BME. BME domes were submerged in type 2 organoid medium [base medium with 20% Wnt3a conditioned medium ([9], H. Clevers lab, Hubrecht Institute), 10% R-spondin 1 conditioned medium ([9], C. Kuo lab, Stanford University), 10% Noggin conditioned medium ([9], H. Clevers lab, Hubrecht Institute), 1x B27 (Gibco, 17504044), 10 μM Nicotinamide (Sigma-Aldrich, N0636), 1.25 mM N-acetylcysteine (Sigma-Aldrich, A0737), 500 ng/ml Hydrocortisone (Sigma-Aldrich, H0888), 100 nM β-estradiol (Sigma-Aldrich, E4389), 10 μM forskolin (Sigma-Aldrich, F6886), 5 μM Y-27632, 5 μM heregulin β1 (Peprotech, 100-03), 20 ng/ml FGF-10 (Peprotech, 100-26), 500 nM A83-01 (Sigma-Aldrich, SML0788), and 5 ng/ml EGF (Peprotech, AF-100-15)]. Organoid cultures were kept at 37 °C and 5% CO_2,_ and media was refreshed every 2-3 days. For passaging, organoid domes were digested with TrypLE Express (Gibco, 12604013) for 2 min at 37 °C. After three washes with Advanced DMEM/F12, organoids were centrifuged at 300 x g for 3 min and the pellet was resuspended in BME for re-plating.

### Organoid histology

For histologic evaluation, organoid domes were embedded in HistoGel (epredia, HG-4000-012) and fixed for 24 h in 10% formalin. After washing with 70% ethanol, organoids were embedded in paraffin according to standard protocols for sectioning. Organoid sections were de-paraffinized and re-hydrated by a xylene/ethanol dilution series. Organoid sections were stained with hematoxylin (Sigma-Aldrich, MHS32) for 4 min and 1% Eosin Y (Sigma-Aldrich, 1.17081.1000) for 10 min according to standard protocols. Organoid sections were imaged with an ECHO Revolve microscope.

### Whole organoid immunofluorescence staining

Whole organoid staining was performed as described previously. [13] Briefly, organoids were released from the BME matrix by incubation with Cell Recovery Solution (Gibco, 354253) for 30-60 min at 60 rpm and 4 °C. After washing with PBS, organoids were fixed with 4% paraformaldehyde (Thermo Scientific, 047392.9M) for 45 min at 4 °C. Organoids were then blocked in 2% bovine serum albumin in PBS with 0.1% Triton-X100 for 15 min at 4 °C. Organoids were incubated with primary antibodies in blocking buffer for 48 h at 4 °C. The following antibodies were used: Keratin 14 conjugated to AF488, clone LL002 (Novus Biologicals, NBP2-34675AF488) at 1:100 dilution, and Keratin 19 conjugated to AF555, clone EP1580Y (abcam, ab203444) at 1:100 dilution. After washing, organoids were incubated with 4’,6-diamidino-2-phenylindole, dilactate (DAPI, Invitrogen, D3571) at a 1:3,000 dilution for 30 min at room temperature. Organoids were cleared by incubation with a 60% glycerol/2.5 M fructose solution for 20 min at room temperature. Organoids were imaged with a Leica SP8 confocal microscope.

### Flow cytometry

Organoids were digested to single cells by digestion with TrypLE Express at 37 °C for 10 min. Cells were blocked with 10% goat serum for 15 min at 4 °C, followed by incubation with antibodies for 45 min at 4 °C. The following antibodies were used: CD326 (EpCAM) conjugated to AF647, clone 9C4 (BioLegend 324212) and CD271 (NGFR) conjugated to FITC, clone Me20.4 (BioLegend, 345104). As a control, cells were incubated without antibodies. Cells were then analyzed using an Attune Flow Cytometer (Thermo Fisher).

### Statistical analyses

For categorical variables, a two proportion Z-test was used to calculate statistical significance. For quantitative variables, a two-tailed Student’s t-test was used to calculate statistical significance. Statistical significance threshold was defined as p-value of less than 0.05.

## Results

CNB was performed on 18 breast excision specimens, and FNAB was performed on 11 breast excision specimens from patients with biopsy-proven DCIS (Figures 1–2). Tumor cells were identified on 16/18 touch preps for CNB specimens and 7/11 direct smears for FNAB specimens (Table 1 and Figure 3). The cytomorphology of the FNAB-sampled DCIS correlated closely with the final tissue histomorphology (Figure 3). CNB-sampled specimens and FNAB-sampled specimens resulted in successful PDO cultures in the majority of cases: 15/18 for CNB samples and 7/11 for FNAB samples (Table 1, Figure 4). Remarkably, the time to initial signs of organoid growth was 19.3 days for CNB-sampled specimens compared to just 4 days for FNAB-sampled specimens (Table 1, Figure 4).

**Table 1.** Rate of PDO generation from FNAB vs. CNB sampled specimens.

|  | **FNA biopsy** | **Core needle biopsy** | **P-value** |
| --- | --- | --- | --- |
| **Tumor cells present on direct smear (for FNAB) or touch prep (for CNB)** | 7/11 (64%) | 16/18 (89%) | 0.10 |
| **Successfully generated patient-derived organoid** | 7/11 (64%) | 15/18 (83%) | 0.23 |
| **Time to initial organoid growth (days)** | 4 (range 1-13) | 19.3 (range 7-55) | 0.01* |

\*p<0.05, two-tailed Students' t-test for quantitative variables, two proportion Z-test for categorical variables

**Figure 3.**
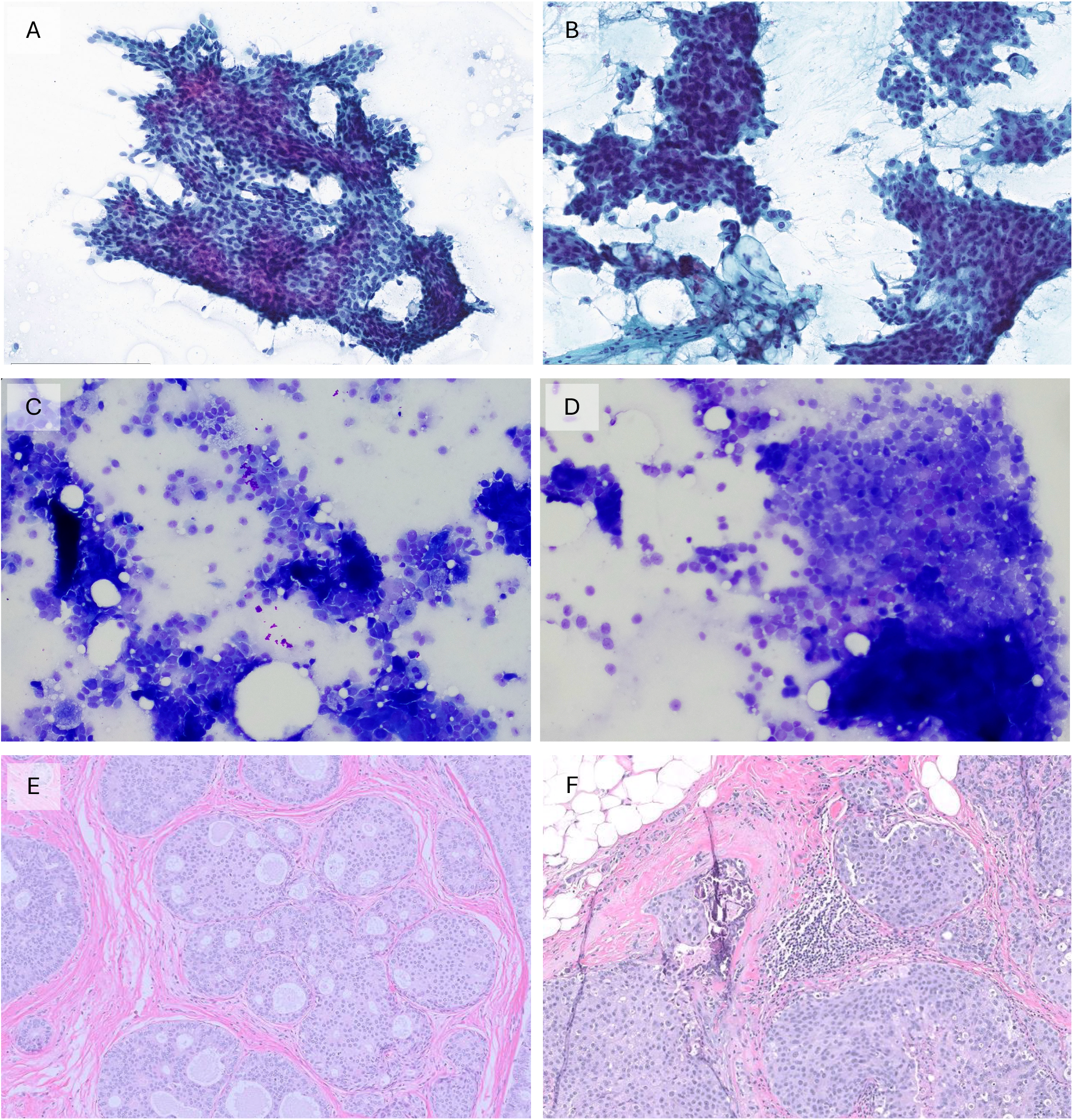
Direct smears, touch preps, and corresponding histology. (A and B) Representative images of direct smears from FNAB. (C and D) Representative images of touch preps from CNB. (E) Representative image of histologic correlate to FNA direct smear in (A). (F) Representative image of histologic correlate to FNA direct smear in (B).

**Figure 4.**
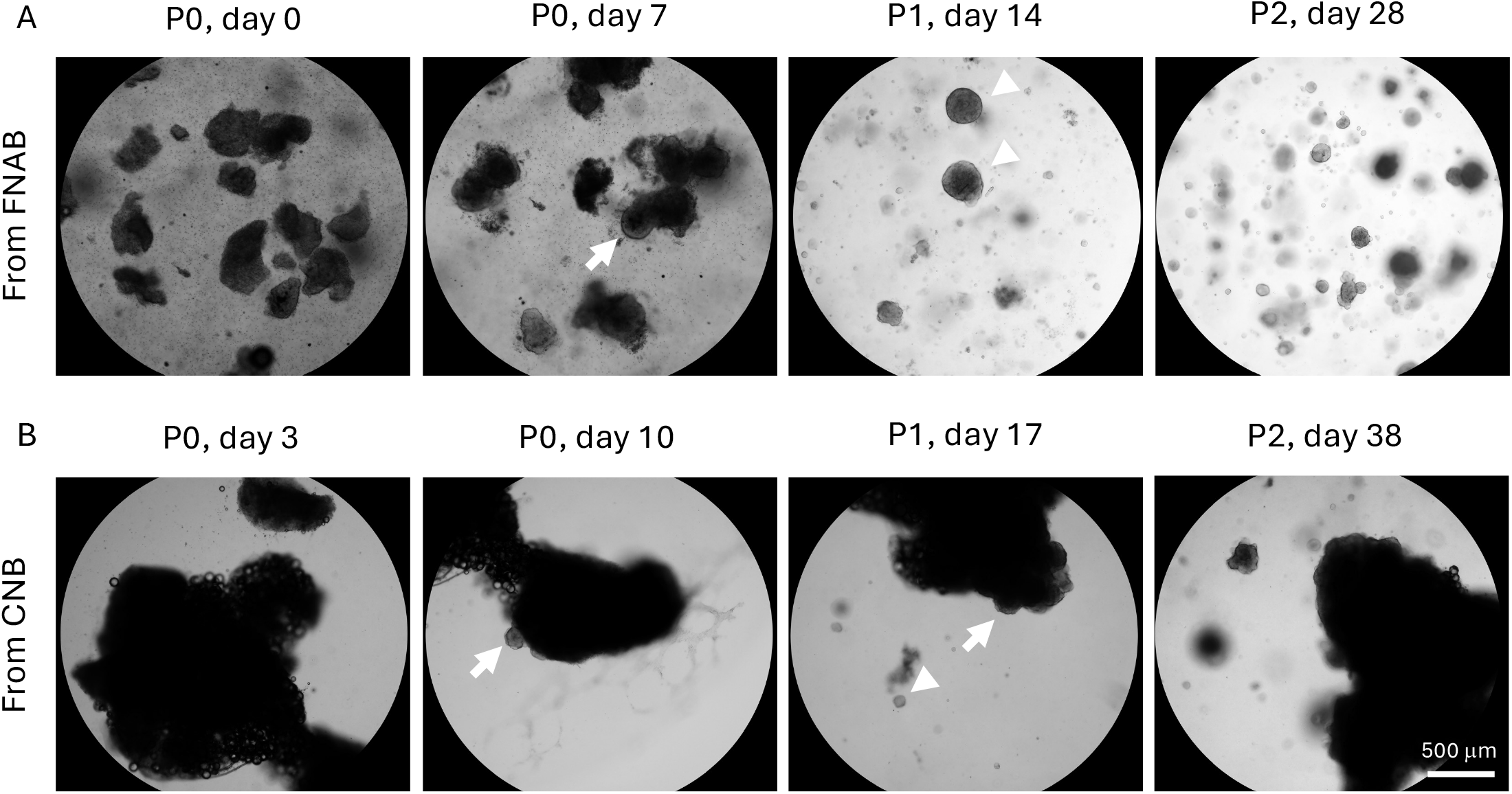
Representative images of DCIS organoid cultures. (A) Organoid culture derived from specimen sampled by FNAB. (B) Organoid culture derived from specimen sampled by CNB. P indicates passage number. Arrows indicate organoids budding off tissue fragments. Arrowheads indicate established organoids.

PDOs were able to be propagated over multiple passages up to 13 passages or 220 days in culture (Figure 4) and demonstrated cribriform growth pattern on histology (Figure 5A), as well as retained myoepithelial cells and luminal cells by immunofluorescence staining and flow cytometry (Figure 5B and C).

**Figure 5.**
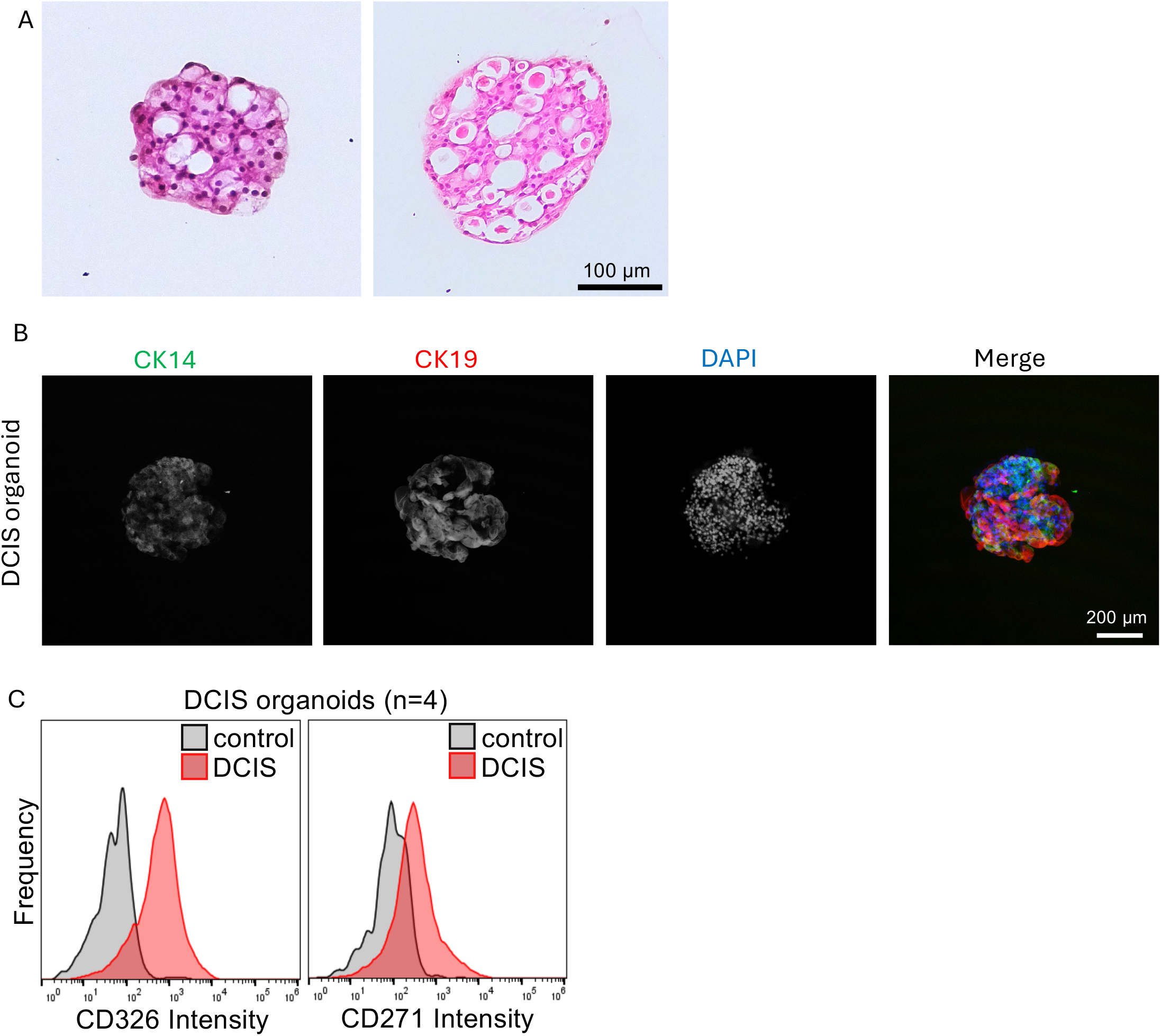
Characterization of DCIS organoids. (A) Representative H&E images of DCIS organoids. (B) Representative confocal images of a DCIS organoid stained with CK14, CK19, and DAPI nuclear stain. DCIS organoid shown was derived from a specimen sampled by FNAB. (C) Histogram of CD326 (EPCAM) and CD271 (NGFR) expression in DCIS organoids analyzed by flow cytometry. Control represents DCIS organoid cells without antibody staining. n = 4 organoids.

Evaluation of the final pathological features of the sampled cases demonstrated that most specimens consisted of intermediate-to-high nuclear grade DCIS that was positive for estrogen receptor (Table 2). Invasive carcinoma was present in CNB-sampled and FNAB-sampled cases at similar frequencies: 4/11 for CNB-sampled cases and 6/18 for FNAB-sampled cases (Table 2). Additionally, among cases that successfully yielded PDOs, invasive carcinoma was also present at similar frequencies for CNB-sampled and FNAB-sampled cases: 6/15 for CNB-sampled cases and 3/7 for FNAB-sampled cases (Table 2). There was no statistically significant correlation between DCIS nuclear grade, biomarker status, and presence of invasive carcinoma on the rate of PDO generation or the time to initial organoid growth (Table 3).

**Table 2.** Final pathological features of specimens sampled by FNAB vs. CB.

|  | **FNA biopsy** | **Core needle biopsy** |
| --- | --- | --- |
| **DCIS nuclear grade** |  |  |
| **1-2** | 2/11 (18%) | 1/18 (6%) |
| **2** | 3/11 (27%) | 7/18 (39%) |
| **2-3** | 2/11 (18%) | 3/18 (17%) |
| **3** | 4/11(36%) | 7/18 (39%) |
| **ER+ DCIS** | 9/11 (82%) | 14/18 (78%) |
| **Invasive carcinoma present in final pathology** | 4/11 (36%) | 6/18 (33%) |
| **Invasive carcinoma present in final pathology**<br>**for specimens with successful organoid**<br>**generation** | 3/7 (43%) | 6/15 (40%) |

**Table 3.** Correlation of final pathologic features with PDO generation.

|  | **Successfully generated**<br>**patient-derived organoid** | **Time to initial organoid**<br>**growth (days)** |
| --- | --- | --- |
| **DCIS nuclear grade**<br>1-2, 2<br>2-3, 3 | 11/13 (85%)<br>11/16 (69%)<br>p = 0.32 | 13.5 (range 2-41)<br>15.4 (range 1-55)<br>p = 0.76 |
| **Biomarker status**<br>ER+<br>ER- | 18/23 (78%)<br>4/6 (67%)<br>p = 0.55 | 13 (range 2-41)<br>20.8 (range 1-55)<br>p = 0.33 |
| **Invasive carcinoma present**<br>Yes<br>No | 9/10 (90%)<br>13/19 (68%)<br>p = 0.2 | 12.3 (range 1-31)<br>15.8 (range 2-55)<br>p = 0.57 |

## Conclusions

Our study demonstrates that FNAB is an effective and efficient method for extracting cells for PDOs modeling DCIS. We were able to achieve high success rates with both culture methods. However, FNAB-sampled specimens yielded visible PDOs significantly more quickly than their

CNB-sampled counterparts, possibly due to the gentler and more complete tissue dissociation afforded by the shear forces of a small-gauge needle, as compared to the mechanical mincing that was utilized for CNB specimens in this study. Our finding is significant for the organoid research community, given the labor-intensive nature of establishing and maintaining organoid cultures, as well as the high cost of the involved reagents.

Although others have described FNAB as a method for obtaining material for PDOs from various invasive tumors, [11,12] our study is unique, as we demonstrate the feasibility of FNAB for sampling breast tumors, and specifically of a pre-invasive lesion. In addition, ours is the first study to compare the efficacy of different biopsy techniques for tissue acquisition. Finally, we demonstrate that the resulting DCIS organoids retain both luminal and myoepithelial cell lineages. This finding is significant as 2D culture of human breast tissue often leads to loss of cellular heterogeneity with a resulting overrepresentation of a single epithelial lineage. [14,15]

The limitations of our study are that we did not compare the efficacy of CNB sampling with enzymatic digestion, which is a widely used processing method for organoid generation, because our experience is that enzymatic digestion may decrease the success rate of organoid culture establishment. In addition, we did not specifically assess for the presence or absence of stromal cells in organoid cultures. However, visual inspection of the culture components suggests that some stromal cell types such as adipocytes and fibroblasts may be present in early passage CNB-derived organoid cultures to a greater extent than in FNAB-derived organoid cultures (data not shown). These observations are consistent with those of prior studies demonstrating that FNAB tends to preferentially capture cellular elements, particularly neoplastic epithelial cells, over acellular stromal components. [16] Finally, we did not quantify the volume of invasive carcinoma present in the excision specimens, leaving open the possibility that the organoid outgrowth could be influenced by the presence of invasive carcinoma with an extensive intraductal component. However, the fact that invasive carcinoma was present in the CNB-sampled and FNAB-sampled specimens at comparable rates (Table 2) suggests against volume of invasive carcinoma as being a factor in the difference in the time to organoid growth.

Taken together, FNAB offers a promising alternative to CNB for breast PDO culture, because it reduces the time and resources that are spent on establishing PDO cultures.

## Data availability statement

The data that support the findings of this study are available from the corresponding author upon reasonable request.

## Author contributions statement

JY and NG conceptualized the study, performed data curation, formal analysis, writing-original draft, and writing-review and editing. RB and JMR conceptualized the study, performed data curation, formal analysis, and writing-review and editing. PV, SW, GCDC, CMR, MMM, KR, SA, MA, CE, KG, RM, JW, and LE performed data curation and writing-review and editing.

## Acknowledgments

We thank the UCSF surgical staff, Agnes Chan and the UCSF Pathology gross room staff, and the UCSF Breast Oncology Program interns for assistance with specimen acquisition and transport. We thank Calvin Kuo and Hans Clevers for providing reagents.

